# Approach for Semi-Automated Measurement of Fiber Diameter in Murine and Canine Skeletal Muscle

**DOI:** 10.1101/569780

**Authors:** Courtney R. Stevens, Michael Sledziona, Timothy P. Moore, Lynn Dong, Jonathan Cheetham

**Affiliations:** Cornell College of Veterinary Medicine, Cornell University, Ithaca, NY

**Author notes:** Address Correspondence to: Jonathan Cheetham, VetMB DipACVS PhD., Department of Clinical Sciences College of Veterinary Medicine Cornell University, Ithaca, NY 14853.

**Keywords:** Feret Diameter, Collagen V, Murine, Canine

## Abstract

Currently available software tools for automated segmentation and analysis of muscle cross-section images often perform poorly in cases of weak or non-uniform staining conditions. To address these issues, our group has developed the MyoSAT (Myofiber Segmentation and Analysis Tool) image-processing pipeline.

MyoSAT combines several unconventional approaches including advanced background leveling, Perona-Malik anisotropic diffusion filtering, and Steger’s line detection algorithm to aid in pre-processing and enhancement of the muscle image. Final segmentation is based upon marker-based watershed segmentation.

Validation tests using collagen V labeled murine and canine muscle tissue demonstrate that MyoSAT can determine mean muscle fiber diameter with an average accuracy of ~97%. The software has been tested to work on full muscle cross-sections and works well even under non-optimal staining conditions.

The MyoSAT software tool has been implemented as a macro for the freely available ImageJ software platform. This new segmentation tool allows scientists to efficiently analyze large muscle cross-sections for use in research studies and diagnostics.

## Introduction

Skeletal muscle is an adaptive tissue, which undergoes changes in mass and fiber composition in response to a wide range of stimuli including exercise, aging, trauma, as well as myopathic and neurological disease. Changes in muscle mass are primarily observed to be associated with atrophy or hypertrophy of individual myofibers as opposed to changes in fiber number. (1,2) Thus, characterization of fiber size distribution in the muscle tissue has significant diagnostic importance.

Muscle fiber size is routinely determined through imaging and analysis of fixed or frozen cross-sections. The fiber size distribution is typically quantified in term of cross-sectional area (CSA) or fiber diameter. The use of minimum feret diameter is preferred as it is the least affected by distortion due to oblique cross-sectioning of muscle tissue. (3)

The development of fluorescent immunohistochemistry (IHC) protocols which label the muscle fiber plasma membrane or extracellular matrix enable high contrast imaging of the fiber boundaries. Effective staining protocols for delineating muscle fibers include dystrophin (4), laminin (5), or collagen (6) staining techniques.

Despite availability of these labelling procedures to aid in identification of the fiber boundaries, segmentation and analysis of scans of muscle cross-sections is still most often accomplished using manual techniques. This is frequently done using basic image annotation software combined with a graphic tablet or mouse. This manual quantification process is tedious and time consuming (7). Considerable regional variability in fiber size is often observed across a muscle section and so a large number of regions must be sampled across the specimen to accurately quantify the fiber size distribution in the overall muscle.

In attempts to speed up this process, several groups have described image-processing frameworks for the automatic segmentation and analysis of muscle cross-sections.

A typical image processing pipeline for muscle cross-sections requires several steps including pre-processing, segmentation, and morphological analysis.(8) The pre-processing step often involves re-adjusting intensity and contrast, background suppression, as well as to providing noise and artifact reduction of the original image. The segmentation step attempts to separate the muscle fibers from background. Finally, morphological analysis extracts feature data such as the fiber size distribution from the segmented image.

Common approaches for automated segmentation of muscle fiber cross-sections range from simple thresholding based strategies (7) to more advanced methods including active contour (9–13) and watershed based algorithms (6,14–17). Unfortunately, while a number of image processing approaches to muscle cross-section analysis have been described in the literature, to date, only a limited number of research groups (13,14,16) have made their computer code available for general use by the research community.

In our experience, a common limitation of currently available software for automated analysis of muscle cross-sections is that segmentation accuracy tends to be highly sensitive to the quality of the input images. Technical issues include weak staining contrast, regional variations in intensity, non-specific staining, as well as presence of artifacts associated with freezing or sectioning (11). These issues can result in mis-segmentation, which may require extensive manual correction. Because currently available software tools are so sensitive to these imaging conditions, their use has not yet gained broad acceptance as a practical tool for research studies and diagnostics.

In this paper, we present MyoSAT (Myofiber Segmentation and Analysis Tool), a semi-automated image processing pipeline that our group has developed to allow analysis of large (and even entire) muscle cross section images. Our goal in developing the MyoSAT software is to offer an improved method, which performs well even in cases of non-ideal staining conditions. The capability to analyze large regions of muscle enables the researcher to identify subtle changes in fiber size distribution between treatment groups.

## Materials and Methods

### Tissue Samples

Animal tissues used for development of the image analysis method were obtained as part of ongoing research studies associated with peripheral nerve repair. No animals were sacrificed specifically for purposes of this study.

Ethics Statement: All associated research studies were performed in accordance with the PHS Policy on Humane Care and Use of Laboratory Animals, NIH Guide for Care and Use of Laboratory Animals, federal and state regulations, and was approved by the Cornell University Institutional Animal Care and Use Committee (IACUC, protocol #2012-0099).

Murine *tibialis-anterior* (TA) muscle tissue: Six C57 BL/6 mice (three male and three female) underwent a left side proximal-tibial to distal-common peroneal cross suture surgery with a conduit repair (18). Five weeks after injury, animals were euthanized and bi-lateral TA muscle cross sections were obtained.

Canine *crico-arytenoid lateralis* (CAL) muscle tissue: Five CAL muscle cross sections were obtained from each of two female beagle dogs with no history of upper airway disease and normal laryngeal function, which was determined endoscopically.

### Immunohistochemistry

The TA and CAL muscle sections were briefly fixed in cold acetone for 10 minutes. 8 µm cryosections were then obtained. The sections were washed in phosphate buffered saline containing 0.05% Tween 20 (PBST) and incubated with 10% rabbitserum and then goat anti-type V collagen antibody (Southern Biotech, Birmingham, AL) at 1:1000 for 1.5 hours. The sections were then further incubated with biotinylated rabbit anti-goat IgG (Vector Laboratories, Burlingame, CA) and streptavidin-Texas Red (Molecular Probe, Life Technologies, Grand Island, NY) to visualize staining. Finally, the sections were stained with DAPI and mounted in ProLong® Diamond anti-fade mountant. As a negative control, goat IgG was diluted to the same final concentration as the primary antibody.

### Microscopy

The Murine TA cross sections were imaged using a Leica Aperio FL slide scanner at 20x. Image resolution: 0.462 µm/pixel (10bits/pixel). The images were saved in 16 bit TIFF format.

Canine CAL cross sections were imaged using an Olympus AX 70 compound fluorescence microscope at 20x with an Optronics MicroFire camera. Original image resolutions: 0.368 μm/pixel (12 bits/pixel). Acquired images were re-scaled using ImageJ to match the 0.462 μm/pixel resolution used for the Murine scans above and re-saved in TIFF format.

### Image Processing Development

The MyoSAT image-processing pipeline consists of 5 stages: 1) Intensity Leveling. 2) Contrast Enhancement. 3) Ridge Detection. 4) Ridge Image Post-Processing. 5) Watershed Segmentation.

#### Intensity Leveling

In typical IHC stained muscle cross-sections, regional fluorescent intensity of the interiors of the muscle fibers is observed to vary across the sample due to non-specific staining as well as variations in tissue thickness. A background leveling technique is applied to suppress this variation. To accomplish this, first the background intensity of the fiber interiors is estimated by applying a median filter with kernel size of 86×86 pixels (40×40 µm). Next, pixel values of the original image (Fig. 1A) are divided by the median filtered image values, which results in a new “leveled” image with each pixel represented with a floating point value. Fiber interiors in this leveled image have a normalized average intensity of ~1.0 and the stained fiber boundaries have intensity >1.0.

**Figure 1:**
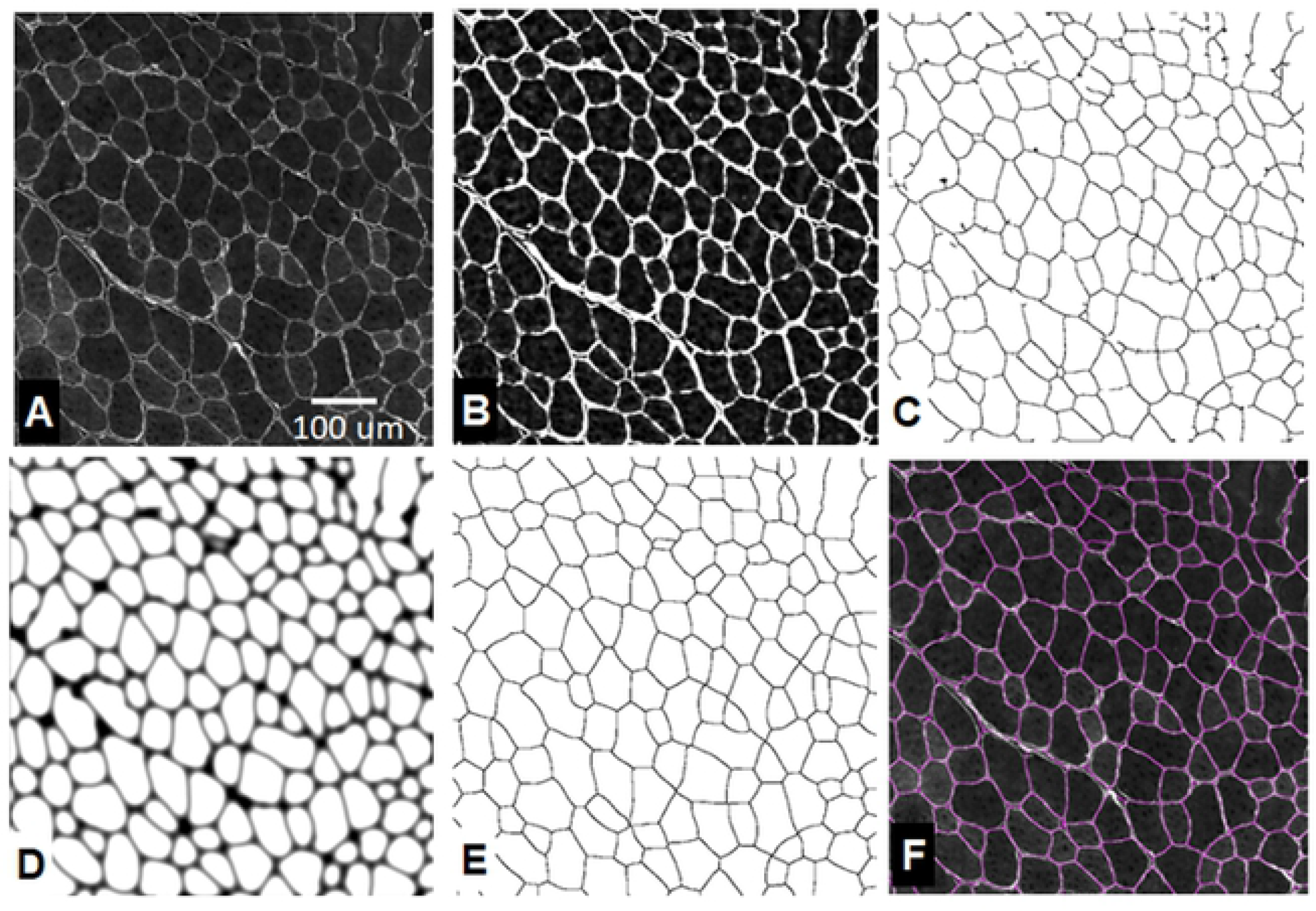
MyoSAT image processing steps: A) Original image. B) Intensity leveling and contrast enhancement. C) Ridge detection. D) Ridge image post-processing. [dilation + Gaussian blur + seed generation] E) Watershed segmentation. F) Result overlay

#### Contrast Enhancement

In the second stage, a Perona-Malik (PM) anisotropic diffusion filter (19) is applied to the new “leveled” image. This filter aids to suppress local image variations and pixel noise while preserving contrast of the fiber boundaries and to enhance the edges. After the PM filter step, the image pixel values are raised to the 4^th^ power which boosts the contrast of the extracellular membranes with respect to background. (Fig. 1B)

#### Ridge Detection

In the third stage, Steger’s line detection algorithm (20) is applied to locate the extracellular membranes between the muscle fibers. An appropriate scale factor (σ) is chosen to maximize detection of the fiber boundaries while rejecting image artifacts. The scale factor is related to the target boundary line width (*w*) by:

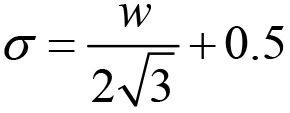

Optimum value of σ was found to range between 4.0 to 8.0 which corresponds to a target line width of between 6 to 12 µm. Output of the ridge detection algorithm is a binary “skeletonized” image containing the detected fiber boundaries. (Fig. 1C)

#### Ridge Image Post-Processing

The output image of the ridge detection algorithm often contains discontinuities as well as detected background artifacts. In order to convert the initial ridge detection image to a final segmented result, several additional post-processing steps are applied: First, a morphological dilation filter (21) using a disk shaped structuring element (radius = 10 pixels) is applied to thicken the detected lines. The next step applies a gaussian blur filter to the binary image. In the final step, a manually adjusted threshold is applied to the blurred image such that values below the threshold are set to zero. This step establishes seed locations for the watershed segmentation algorithm, which is described next. (Fig. 1D)

#### Watershed Segmentation

The final stage of the pipeline applies a classic watershed segmentation algorithm (22) to the blurred image. Regions of zero intensity established by the manually adjusted thresholding step above provide seed locations for the segmentation algorithm. After watershed segmentation (Fig 1E,F), the detected objects are classified by cross-sectional area (CSA) and feret diameter. To aid in filtering out mis-segmented regions, objects with CSA outside a predetermined range (~200 to 10,000 μm^2^) are excluded from the analysis. This range corresponding to the smallest and largest fiber sizes observed in the target muscle tissues.

The image-processing pipeline has been implemented as a macro for the freely available image processing package FIJI / ImageJ [20]. The macro requires several third-party ImageJ plugins which implement several of the image processing algorithms (23) (24) (25).

### Validation

#### Validation Image Sets

The validation image sets consisted of 12 murine TA muscle cross-section images (ROI area: 0.17 – 0.86 mm^2^) and 10 canine CAL muscle cross-section images (ROI area: 0.26mm^2^)

#### Manual Segmentation (Ground Truth)

Fiber boundaries within the muscle cross-section images were manually segmented with the aid of a graphics tablet (Wacom Cintiq 22HD). Feret diameters of the manually segmented regions were then obtained using the ImageJ “Analyze Particles” function.

#### SMASH Software

To compare performance of the MyoSAT with other currently available open-source software for automated muscle cross-section image segmentation, the image sets were similarly segmented using the recently released SMASH (Semiautomatic Image Processing of Skeletal Muscle Histology) software package (16).

#### Accuracy Analysis

Histograms of fiber diameter were generated for each muscle section image using each of the three segmentation methods (manual, MyoSAT, and SMASH). The accuracy of the automated approaches was compared to ground truth by statistical comparisons of mean fiber diameter obtained for each image. For reasons described in the previous section, segmented objects with CSA outside of the pre-determined given range (~200 to 10,000 μm^2^) were excluded from the analysis. Fibers intersecting the ROI edges were also excluded from analysis.

### Development of a Staining Contrast Mapping Tool

It was observed that several of the image-processing pipeline steps could be combined to provide a staining contrast mapping tool valuable for optimization of staining protocols. We define the staining contrast in the image as the intensity ratio between the staining of the fiber boundaries (extracellular membranes) to the non-specific staining of the fiber interiors.

The accuracy of most segmentation algorithms is often highly sensitive to staining contrast within the image. Such contrast information aids in quality control by identifying cross-section regions with sufficient contrast to provide good segmentation accuracy.

The three steps used to generate the staining contrast map are: 1) The original image is divided by the median filtered image to generate a “leveled image”. 2) The ridge detection image generated using Steger’s line detection algorithm is used as a mask to sample the centers of the fiber boundaries to generate the local contrast map. 3) An averaging function with kernel size (27 × 27 µm^2) is then applied to the map to reduce localized intensity variations. This aids in assessing overall staining contrast within a muscle region.

## Results

### Validation Testing

The validation images were analyzed using three different segmentation methods: 1) Manual tracing (ground truth), 2) the proposed MyoSAT image processing method, and 3) the SMASH software package.

The image sets consisted of 12 murine TA muscle cross-section images containing 91 – 165 fibers (mean= 124.9); and 10 canine CAL muscle cross-section images containing 28-153 intact fibers per image (mean=89.3)).

We found that for both murine and canine tissue samples, fiber counts and fiber diameter obtained using the proposed semi-automated muscle segmentation method (MyoSAT) closely correlated to results obtained using manual segmentation of the muscle fibers (Table 1).

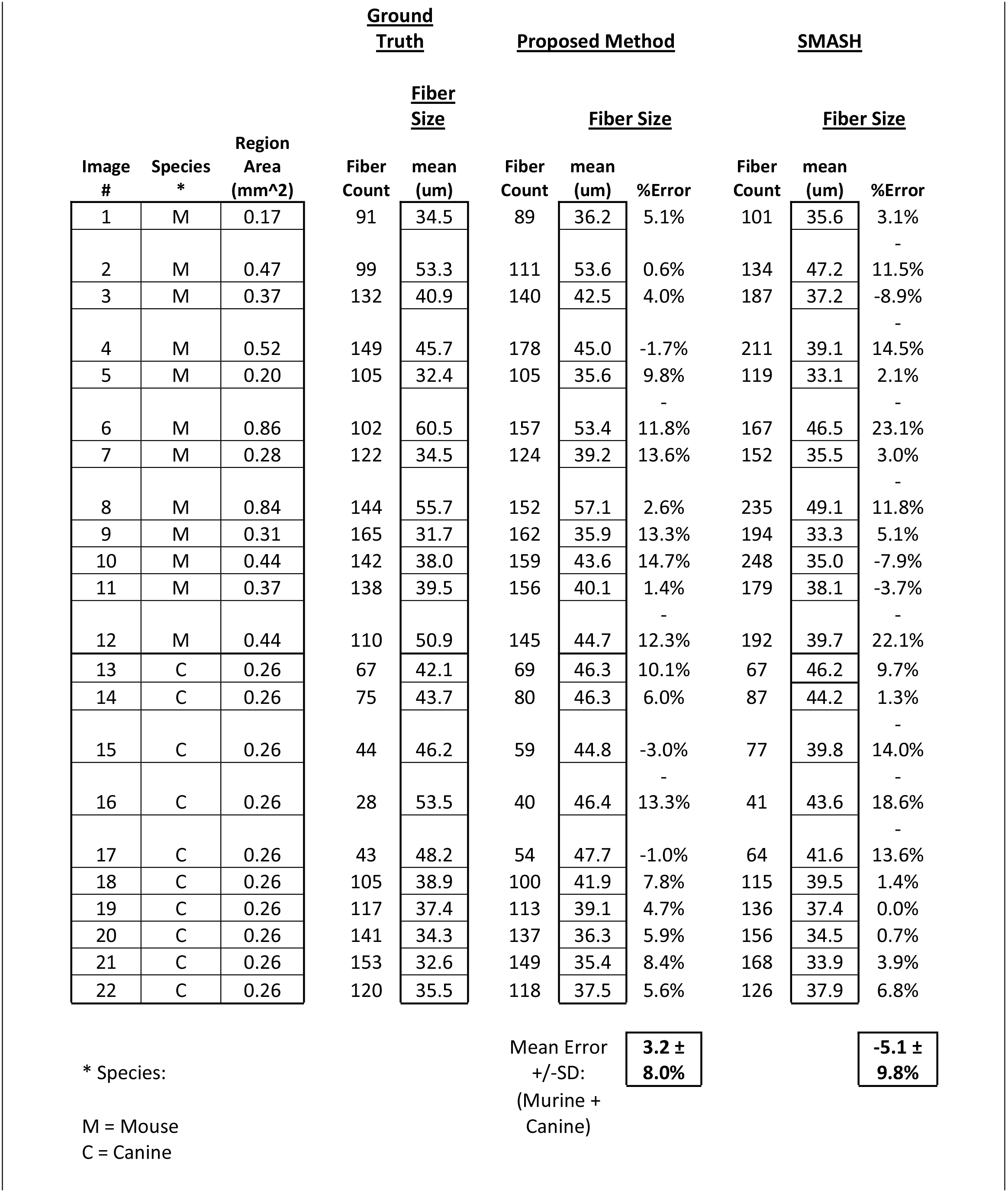
Comparison of Mean Fiber Size Measurement Accuracy.

For the murine TA samples, no significant differences were observed between mean fiber estimates for the proposed method and ground truth (mean difference ± std error, 0.76 ± 1.15µm, p=0.52, paired two-tailed test). Significant differences were observed between ground truth and the SMASH method (4.02 ± 1.46µm, p=0.019, paired two-tailed test).

For the canine CAL samples, no significant differences were observed between mean fiber estimates for the proposed method and ground truth (mean difference ± std error, 0.94 ± 1.03µm, p=0.39, paired two-tailed test). Significant differences were observed between ground truth and the SMASH method (2.32 ± 0.62µm, p=0.005, paired two-tailed test).

The average difference between the proposed method and ground truth for the 22 combined murine and canine sample images was 3.2% [SD=±8.0%]. The difference between the SMASH method and ground truth was −5.1% [SD=±9.8%]. We observed that the SMASH software had a tendency to over-segment the fibers resulting in higher fiber counts and underestimation of the fiber diameters.

Bland-Altman analysis (26) indicates that the MyoSAT method correlates slightly more closely (y = −0.32x + 14) with the manual method than SMASH (y = −0.58x + 2, p = 0.067, ANOVA). (See Fig. 2.)

**Figure 2:**
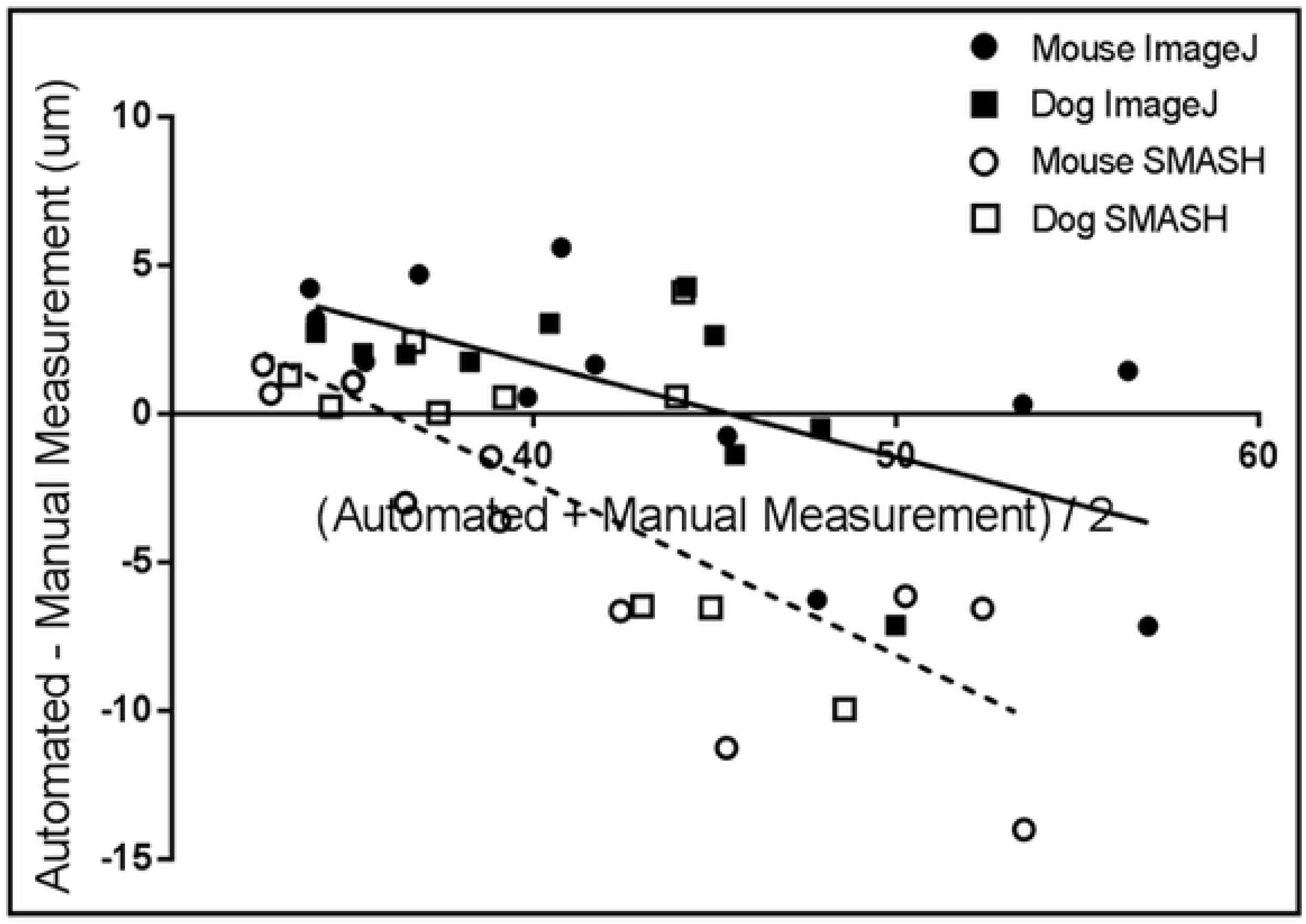
Combined Bland-Altman plot for murine and canine fiber diameter data obtained using MyoSAT, and SMASH compared to manual tracing method (Ground Truth). Linear regression: MyoSAT: y= −0.32x + 14. SMASH: y= −0.58x +21.

We next compared the fiber size histograms generated using manual, MyoSAT, and SMASH segmentation methods. We observed that the MyoSAT analysis pipeline produces fiber size distributions histograms that more closely resemble ones generated by manual segmentation than size distributions generated using the SMASH segmentation approach.

### Analysis of full Muscle Cross-Sections

A primary motivation for the development of the MyoSAT software was to develop a reliable method for segmentation and analysis of large muscle cross-sections regions.

Using the MyoSAT segmentation method to analyze entire murine TA muscle cross-section images frequently identified regions of non-uniform and low contrast staining. Fig. 3 provides an example output of the proposed method. The analysis of the example image took approximately 15 minutes. We found analysis of such large regions to be unreliable using other available image processing software for automatic fiber segmentation due to issues including mis-segmentation, processing time, and image size constraints.

**Figure 3:**
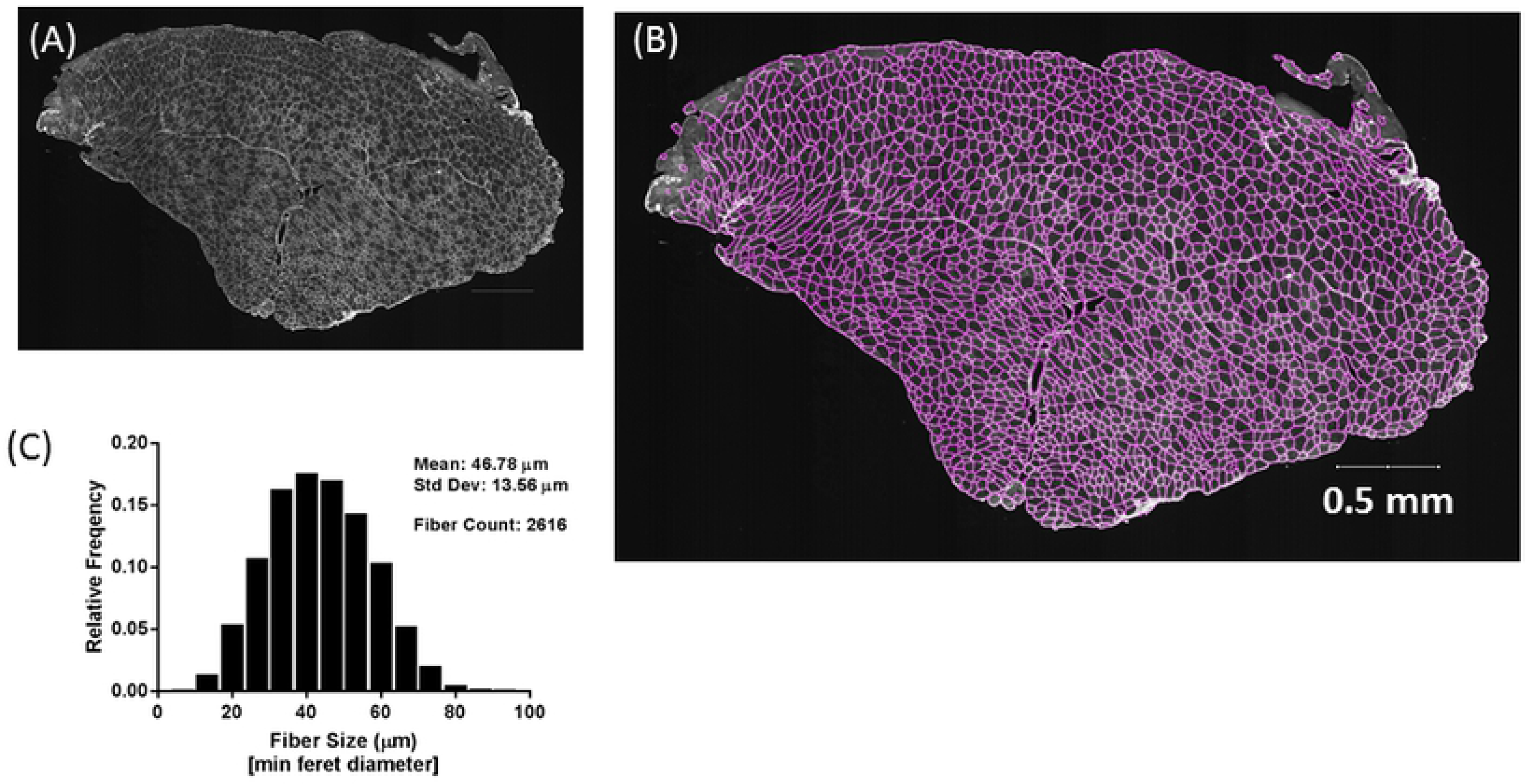
*A,B:*Example of full cross-section of murine TA muscle analyzed using the MyoSAT semi-automatic fiber detection software. *C:* Histogram of (min feret) fiber diameters. MyoSAT has identified n=2616 fibers with mean fiber diameter=46.78µm (SD+/-13.56 µm).

We next demonstrated the application of MyoSAT to identify changes in muscle physiology, Left and right size TA muscle cross-sections were obtained from mice 5 weeks after a unilateral nerve transection and graft. MyoSAT was used to analyze the full TA muscle cross-sections images containing between 1153 - 2637 identified fibers (mean 1784.1).

As anticipated, evidence of reduced muscle fiber diameter was detected in the repaired limb (34.0µm (±0.7µm)) compared to the control (46.4µm (± 2.3µm), figure 4, p=0.0057, n=4 mice, paired t-test). This demonstrates the use of MyoSAT data to quantify and identify significant statistical differences in fiber size obtained via analysis of full-muscle cross-sections. This new method avoids the traditional approach of having to analyze a large number of individual ROI’s of each cross-section.

**Figure 4:**
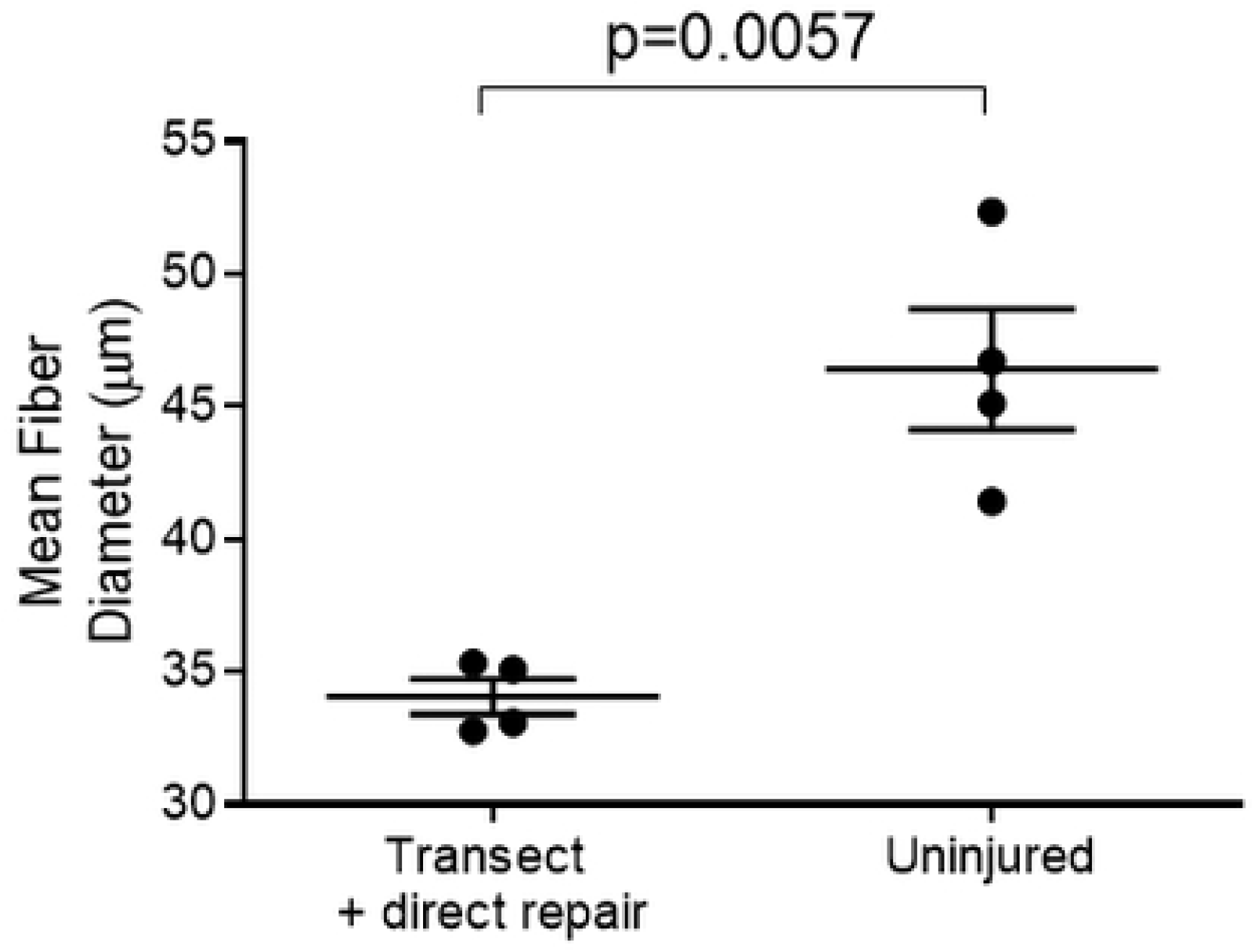
Example MyoSAT analysis used to evaluate change in muscle morphology in murine TA muscle 5 weeks after a left-side tibial-peroneal cross-suture nerve repair. Analysis of the data reveals significant differences in average fiber size between operated (left) and intact (right) sides. (n=4 mice) Error bars=Std. Error.

### Application of Contrast Analysis Tool

We found the staining contrast mapping tool, which we developed as a simple extension of the image-processing pipeline, to be very useful in providing an objective assessment of staining contrast. The contrast map allows the user to quickly identify poorly stained regions, which may result in reduced segmentation accuracy.

Fig. 5 demonstrates output of the contrast-mapping tool applied to a full cross-section of Col V stained murine TA muscle exhibiting a non-uniform staining issue. In this case, the regional average contrast ratio between the fiber boundaries and the fiber interior ranged from approx. 1.0 to 3.0.

**Figure 5:**
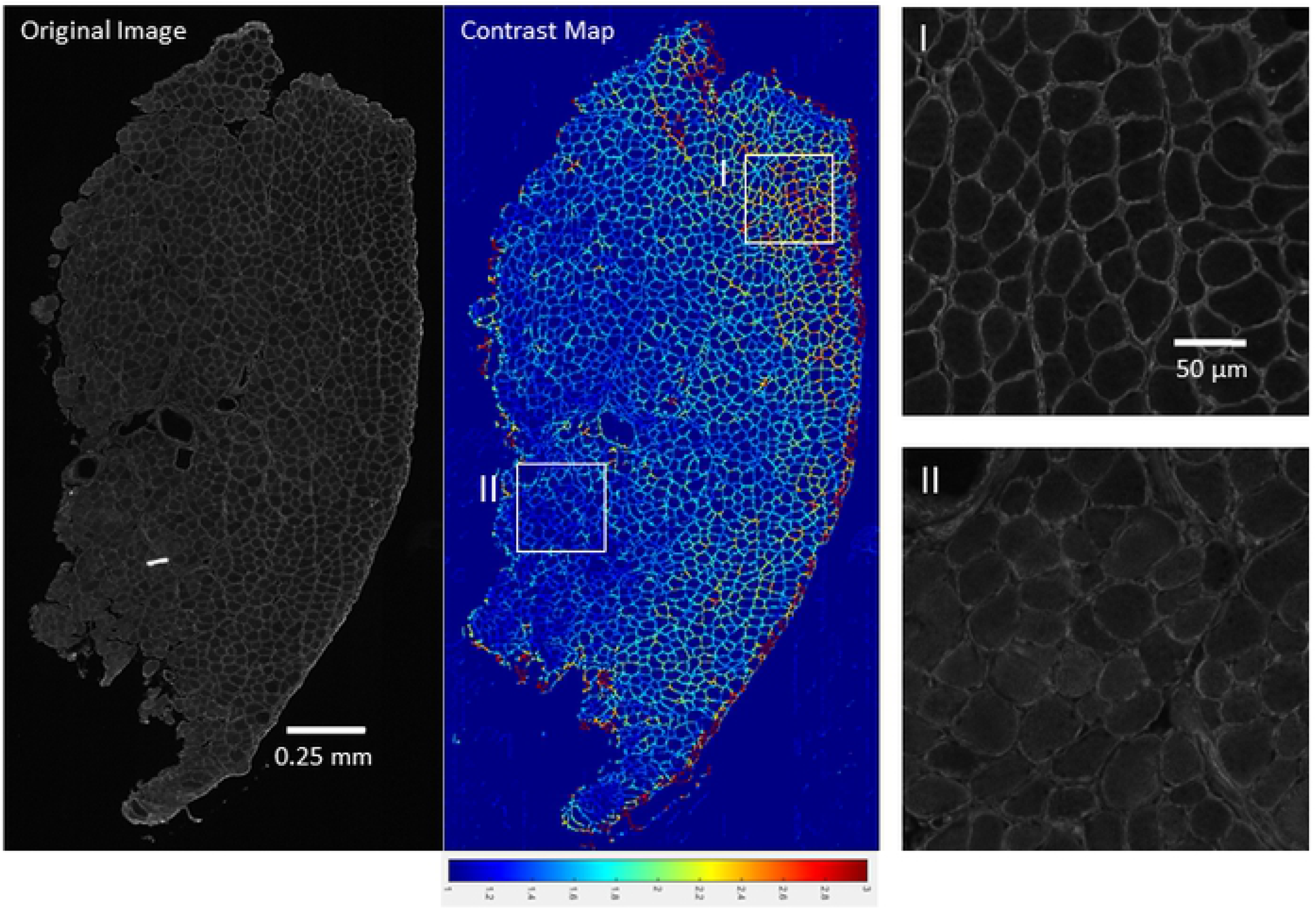
Development of contrast mapping tool to assess staining uniformity and contrast required for accurate fiber segmentation illustrating regions of high contrast (I) vs poor contrast (II) fiber staining. Contrast map range 1.0-3.0.

In our testing, a contrast ratio of approximately >2.25 across a region was found to provide consistent and accurate segmentation of the fiber boundaries using the MyoSAT software.

## Discussion

Here we have introduced a semi-automated image-processing pipeline (MyoSAT) which we developed for accurate segmentation and size distribution analysis of myofibers in muscle cross-section images. As we have described, MyoSAT incorporates several novel pre-processing steps to compensate for non-uniform staining intensity and to enhance the contrast of the fiber boundaries. Several of the unconventional steps within the image-processing pipeline include: 1) Aggressive pre-processing to compensate for non-uniform staining. 2) Application of anisotropic diffusion filtering to provide noise reduction and fiber border enhancement. 3) Use of Steger’s algorithm to detect and binarize the fiber boundaries.

Our validation tests have demonstrated that the MyoSAT analysis provides on average 96% accuracy for estimation of mean fiber diameter when compared to human segmented images in both murine and canine muscle tissue. Muscle fiber size distribution histograms generated during the analysis were found to closely approximate results obtained by manual segmentation.

It is important to note that the MyoSAT segmentation approach is based upon a line detection algorithm to detect the fiber boundaries, whereas most other software approaches are based upon edge detection algorithms. As a result, MyoSAT establishes segmentation boundaries halfway between the individual fibers instead of attempting to trace the fiber interior edges. As such, the proposed method has limited ability to reject the presence of excessive connective tissue in a region. Despite this limitation, the primary advantage of the line detection approach is the enhanced ability to segment low contrast images. This approach also avoids subjectivity associated with identifying the precise location of each fiber membrane edge, which is a common issue with the edge based detection algorithms.

MyoSAT is a semi-automated segmentation approach and so some manual adjustment of sensitivity parameters is required in order to achieve accurate segmentation. A disadvantage to this approach is that manual tuning can introduce some subjectivity into the analysis. However, an advantage is the ability to adjust the image processing to work for a wide range of imaging and staining conditions.

As with most automated image segmentation approaches, the segmentation accuracy of the proposed method is ultimately limited by staining contrast and image quality. A secondary outcome of our work has been the development of a contrast-mapping tool, which we have found to be useful for optimization and quality control of muscle staining protocols. The contrast mapping technique provides an objective approach to identify regions with sufficient staining contrast in order to yield accurate segmentation.

Our research group developed MyoSAT after having limited success with both commercial and open-source software packages to reliably analyze muscle fiber size distributions in large cross-sections, which we needed for own work. MyoSAT can be applied to entire cross sections of skeletal muscle, allowing the accurate segmentation and measurement of thousands of individual fibers.

Future development goals for the MyoSAT software include: 1) Development of a user-friendly interface. 2) Optimization of the image processing algorithms to improve performance. 3) Incorporate additional staining channel data to take advantage of ATPase (27) or myosin heavy chain (28) staining techniques to provide capability for fiber type distribution analysis.

The MyoSAT image-processing pipeline has been implemented as a macro within the freely available image processing package FIJI / ImageJ [20]. The macro may be downloaded from XXX. The availability of this software will enable the research community to efficiently analyze large muscle cross-sections for experimental studies and diagnostics.

## Acknowledgements

None.

## References

1. Taylor NAS, Wilkinson JG. Exercise-Induced Skeletal Muscle Growth Hypertrophy or Hyperplasia? Sport Med An Int J Appl Med Sci Sport Exerc. 1986;3(3):190–200.

2. Nicks DK, Beneke WM, Key RM, Timson BF. Muscle fibre size and number following immobilisation atrophy. J Anat. 1989;163:1–5.

3. Briguet A, Courdier-Fruh I, Foster M, Meier T, Magyar JP. Histological parameters for the quantitative assessment of muscular dystrophy in the mdx-mouse. Neuromuscul Disord. 2004;14(10):675–82.

4. McCarthy JJ, Mula J, Miyazaki M, Erfani R, Garrison K, Farooqui AB, Srikuea R, Lawson BA, Grimes B, Keller C, Van Zant G, Campbell KS, Esser KA, Dupont-Versteegden EE, Peterson CA. Effective fiber hypertrophy in satellite cell-depleted skeletal muscle. Development. 2011;138(17):3657–66.

5. Roorda BD, Hesselink MKC, Schaart G, Moonen-Kornips E, Martínez-Martínez P, Losen M, De Baets MH, Mensink RP, Schrauwen P. DGAT1 overexpression in muscle by in vivo DNA electroporation increases intramyocellular lipid content. J Lipid Res. 2005;46(2):230–6.

6. Sáez A, Rivas E, Montero-Sánchez A, Paradas C, Acha B, Pascual A, Serrano C, Escudero LM. Quantifiable diagnosis of muscular dystrophies and neurogenic atrophies through network analysis. BMC Med. 2013;11(1).

7. Castleman KR, Chui LA, Martin TP, Edgerton VR. Quantitative muscle biopsy analysis. Monogr Clin Cytol. 1984;9(15):101–16.

8. Miazaki M, Viana MP, Yang Z, Comin CH, Wang Y, da F Costa L, Xu X. Automated high-content morphological analysis of muscle fiber histology. Comput Biol Med. Elsevier; 2015;63:28–35.

9. Brox T, Kim Y, Weickert J, Feiden W. Fully-automated Analysis of Muscle Fiber Images with Combined Region and Edge Based Active Contours. 2006;86–90.

10. Klemencic A, Kovacic S, Pernus F. Automated segmentation of muscle fiber images using active contour models. Cytometry. Faculty of Electrical Engineering, University of Ljubljana, Slovenia.; 1998 Aug 1;32(4):317–26.

11. Liu F, Mackey AL, Srikuea R, Esser KA, Yang L. Automated image segmentation of haematoxylin and eosin stained skeletal muscle cross-sections. J Microsc. 2013;252(3):275–85.

12. Mula J, Lee JD, Liu F, Yang L, Peterson CA. Automated image analysis of skeletal muscle fiber cross-sectional area. J Appl Physiol (Bethesda, Md 1985). College of Health Sciences, University of Kentucky, Lexington, Kentucky 40536, USA.; 2013 Jan 1;114(1):148–55.

13. Wen Y, Murach KA, Vechetti IJ, Fry CS, Vickery CD, Peterson CA, McCarthy JJ, Campbell KS. MyoVision: Software for Automated High-Content Analysis of Skeletal Muscle Immunohistochemistry. J Appl Physiol. 2017;jap.00762.2017.

14. Bergmeister KD, Gröger M, Aman M, Willensdorfer A, Manzano-Szalai K, Salminger S, Aszmann OC. Automated muscle fiber type population analysis with ImageJ of whole rat muscles using rapid myosin heavy chain immunohistochemistry. Muscle and Nerve. 2016;54(2):292–9.

15. Meunier B, Picard B, Astruc T, Labas R. Development of image analysis tool for the classification of muscle fibre type using immunohistochemical staining. Histochem Cell Biol [Internet]. 2010 Sep [cited 2013 Aug 4];134(3):307–17.

16. Smith LR, Barton ER. SMASH - semi-automatic muscle analysis using segmentation of histology: A MATLAB application. Skelet Muscle. 2014;4(1).

17. Strange H, Scott I, Zwiggelaar R. Myofibre segmentation in H&E stained adult skeletal muscle images using coherence-enhancing diffusion filtering. BMC Med Imaging. 2014;14(1):1–13.

18. Žygelytė E, Bernard ME, Tomlinson JE, Martin MJ, Terhorst A, Bradford HE, Lundquist SA, Sledzonia M, Cheetham J. RetroDISCO: Clearing technique to improve quantification of retrograde labeled motor neurons of intact mouse spinal cords. J Neurosci Methods. 2016;271.

19. Perona P, Malik J. Scale-space and edge detection using anisotropic diffusion. IEEE Trans Pattern Anal Mach Intell. 1990;12(7):629–39.

20. Steger C. An Unbiased Detector of Curvilinear Structures. Munchen; 1996.

21. Jain AK. Fundamentals of Digital Image Processing. Upper Saddle River, NJ, USA: Prentice-Hall, Inc.; 1989.

22. Soille P, Vincent LM. Determining watersheds in digital pictures via flooding simulations. 1990;(September 1990):240–50.

23. Sacha J. IJ Plugins Toolkit v2.1.0 (ImageJ Plugin). 2017.

24. Wagner T, Hiner M. Ridge Detection v1.4.0 (ImageJ Plugin). 2017.

25. Legland D, Arganda-Carreras I, Andrey P. MorphoLibJ: Integrated library and plugins for mathematical morphology with ImageJ. Bioinformatics. 2016;32(22):3532–4.

26. Bland JM, Altman DG. Statistical methods for assessing agreement between two methods of clinical measurement. Lancet [Internet]. 1986 Feb 8 [cited 2014 Jul 9];1(8476):307–10.

27. Brooke MH, Kaiser KK. Muscle Fiber Types: How Many and What Kind? Arch Neurol. 1970;23(4):369–79.

28. Schiaffino S, Gorza L, Sartore S, Saggin L, Ausoni S, Vianello M, Gundersen K, Lømo T. Three myosin heavy chain isoforms in type 2 skeletal muscle fibres. J Muscle Res Cell Motil. 1989;

